# Integrin-based adhesions promote cell-cell junction remodelling and cytoskeletal rearrangements to drive embryonic wound healing

**DOI:** 10.1101/2023.03.13.532433

**Authors:** Michelle Ly, Clara Schimmer, Ray Hawkins, Katheryn Rothenberg, Rodrigo Fernandez-Gonzalez

## Abstract

Embryos have a remarkable ability to repair wounds rapidly, with no inflammation or scarring. Embryonic wound healing is driven by the collective movement of the surrounding cells to seal the lesion. During embryonic wound closure, the cells adjacent to the wound polarize the cytoskeletal protein actin and the molecular motor non-muscle myosin II, which accumulate at the wound edge forming a supracellular cable around the wound. Adherens junction proteins including E-cadherin are internalized from the interface with the lesion and localize to former tricellular junctions at the wound margin, in a process necessary for cytoskeletal polarity. Using quantitative live microscopy, we found that the cells adjacent to wounds in the *Drosophila* epidermis also polarized Talin, a core component of cell-extracellular matrix (ECM) adhesions that links integrins to the actin cytoskeleton. Integrin knock-down and inhibition of integrin binding delayed wound closure and were associated with a reduction in actin levels around the wound. Additionally, disrupting integrins caused a defect in E-cadherin reinforcement at tricellular junctions along the wound edge, suggesting crosstalk between integrin-based and cadherin-based adhesions. Together, our results show that cell-ECM adhesion contributes to embryonic wound repair and reveal an interplay between cell-cell and cell-ECM adhesion in the collective cell movements that drive rapid wound healing.

## INTRODUCTION

Embryos repair wounds rapidly, with no inflammation or scarring (Rowlatt, 1979; Whitby and Ferguson, 1991). The mechanisms of embryonic wound repair are conserved across species and involve the collective movement of the cells around the wound (Hunter and Fernandez-Gonzalez, 2017). Cell movement and coordination during tissue repair are associated with cytoskeletal polarization. Actin and the molecular motor non-muscle myosin II are asymmetrically distributed in the cells adjacent to the wound, accumulating at the wound edge (Brock et al., 1996; Martin and Lewis, 1992), where they form a contractile cable that coordinates cell movements (Wood et al., 2002; Zulueta-Coarasa and Fernandez-Gonzalez, 2018). Actomyosin cable assembly requires remodelling of the adherens junctions that mediate cell-cell adhesion: the junctional proteins E-cadherin, β-catenin, and α-catenin are internalized from the interface with the wounded cells (Abreu-Blanco et al., 2012; Brock et al., 1996; Hunter et al., 2015; Matsubayashi et al., 2015; Wood et al., 2002; Zulueta-Coarasa et al., 2014), and accumulate at former tricellular junctions along the wound edge (Abreu-Blanco et al., 2012; Wood et al., 2002; Zulueta-Coarasa et al., 2014). Disrupting adherens junction remodelling prevents actomyosin cable assembly and slows down wound repair (Hunter et al., 2015; Matsubayashi et al., 2015).

Cell migration in many systems involves adhesion to and remodelling of the ECM (Charras and Sahai, 2014; Fraley et al., 2010). Cells adhere to the ECM through integrins, transmembrane receptors formed by α and β subunits that link the ECM to the cytoskeleton via intracellular adaptor proteins. The ECM proteins Collagen IV and Laminin contain integrin binding sites and are commonly implicated in cell-ECM adhesion (Hynes and Zhao, 2000). In 2D cell cultures, integrin-based adhesions (IBAs) termed focal adhesions remodel in response to mechanical cues from actomyosin contractility and substrate stiffness (Fraley et al., 2010; Geiger et al., 2009). Similar to embryonic repair, cells in wounded 2D tissue cultures assemble a contractile actomyosin cable around the wound (Bement et al., 1993). In the presence of ECM, the cable relays tension to the substrate through IBAs (Brugués et al., 2014). IBAs *in vivo* share a similar composition with focal adhesions *in vitro* (Goodwin et al., 2016; Gunawan et al., 2019). In addition, cell-cell and cell-ECM adhesions are mechanically linked both in tissue culture and *in vivo* (Bécam et al., 2005; Borghi et al., 2010; Goodwin et al., 2017; Jülich et al., 2015), suggesting that IBAs may contribute to morphogenetic processes that require cell-cell adhesion remodelling.

## RESULTS AND DISCUSSION

### The ECM is remodelled during *Drosophila* embryonic wound repair

To determine if wound healing in the *Drosophila* embryonic epidermis occurs within an ECM, we examined the presence of ECM proteins midway into embryonic development (stages 14-15), when epidermal cells are already differentiated (Payre, 2003). Using immunofluorescence, we found that epidermal cells in the *Drosophila* embryo were surrounded by collagen and laminin (Figure 1A-B). Both Collagen IV and Laminin A displayed a modest, preferential enrichment at the basal side, with 14±8% and 18±8% (mean±standard error of the mean) higher basal than apical accumulations, respectively (Figure 1A’-A’’, B). These results suggest that the *Drosophila* embryonic epidermis is embedded within a multi-protein ECM.

**Figure 1.**
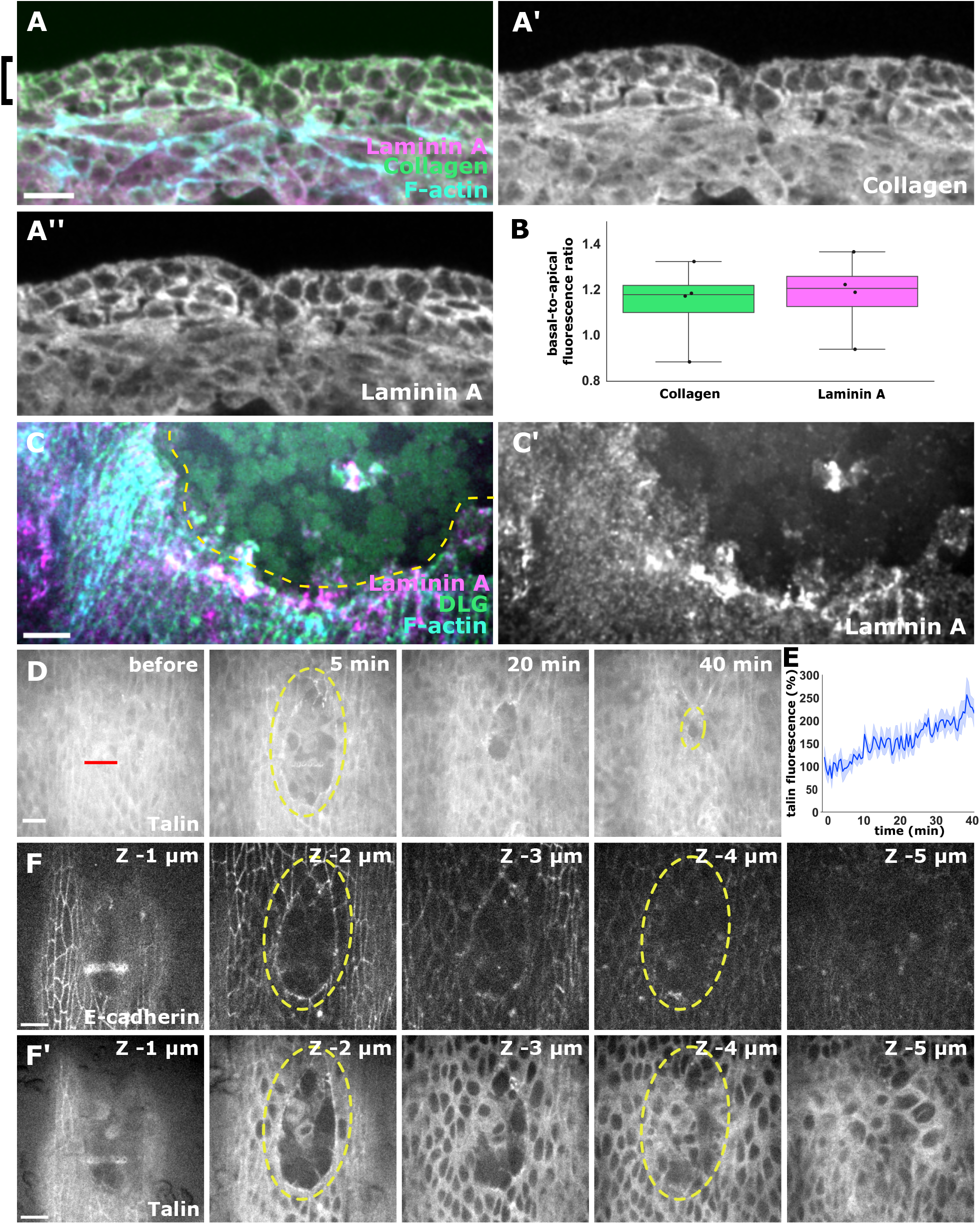
Embryonic wound healing involves cell-ECM adhesion remodelling. **(A, C)** Cross-sections (A) or en-face views (C) of the *Drosophila* epidermis in embryos expressing LamininA:GFP and stained with antibodies against collagen (A’, green in A) or GFP (A’’, C’, magenta in A, C), the basolateral marker Discs Large (DLG, C, green), and with phalloidin to visualize F-actin (cyan in A, C). Black brackets delimit the epidermis. **(B)** Basal-to-apical fluorescence ratio for collagen (green) and Laminin A (magenta) (*n =* 11 basal segments and the corresponding apical segments in 4 embryos). **(D)** Wound healing in an embryo expressing Talin:GFP. Time after wounding is shown. Red line indicates wound site. **(E)** Talin:GFP fluorescence at the wound edge (*n* = 6 wounds). **(F)** Z slices displaying the wound edge 5 minutes after wounding in an embryo expressing E-cadherin:tdTomato (F) and Talin:GFP (F’). (C, D, F) Yellow dashed lines indicate the wound edge. Anterior left (A, C, D, F), dorsal up (C, D, F). Bars, 10 μm.

To determine if wound healing is associated with ECM remodelling, we fixed wounded embryos expressing GFP-tagged Laminin A (Sarov et al., 2016). Laminin accumulated at the wound edge (Figure 1C-C’), suggesting that the ECM is remodeled during wound closure, presumably by IBAs at the wound margin. Consistent with this, a GFP-tagged form of Talin (Yuan et al., 2010), a protein that links IBAs to the actin cytoskeleton (Bulgakova et al., 2012; Goodwin et al., 2016), accumulated at the wound edge by 2.6±0.2 fold over the first 40 minutes of wound healing with respect to the levels immediately after wounding (*P* = 0.03, Figure 1D-E, Movie S1). At the wound edge, talin colocalized with E-cadherin along the apical-basal axis (Figure 1F, Movie S2), suggesting that IBAs and adherens junctions are closely associated at the wound front. *In vitro*, talin is involved in actin nucleation, shortening, and filament crosslinking (Goldmann et al., 1994; Goldmann et al., 1999). Talin is recruited from the cytoplasm by integrins to form the core of the intracellular protein network that connects integrins to the cytoskeleton. Consistent with this functional and mechanical link, talin and integrin mutants share similar phenotypes during *Drosophila* development (Brown et al., 2002). Together, our data suggest that cell-ECM adhesion may contribute to embryonic wound closure.

### Integrins are necessary for rapid embryonic wound healing

To establish if cell-ECM adhesion is necessary for rapid embryonic wound repair, we quantified the dynamics of wound closure in embryos in which integrins had been knocked down by RNA interference (RNAi). The *Drosophila* genome encodes five α integrin subunits, αPS1-5, and two β subunits, βPS and βv (Brown et al., 2000; Hynes and Zhao, 2000). Integrin βPS, encoded by the gene *myospheroid* (*mys*) is the predominant β subunit in the *Drosophila* embryonic epidermis (Brown et al., 2000). We used the UAS-Gal4 system (Brand and Perrimon, 1993) to drive a transgenic RNAi line against *mys* (Perkins et al., 2015). *daughterless-Gal4* (*da-Gal4*)-driven RNAi was expressed throughout the epidermis and reduced Mys levels by 33% (Figure S1A-C, *P* = 0.03), resulting in 100% embryonic lethality. *mys* RNAi embryos repaired wounds 54% slower than *mCherry* RNAi controls (*P* = 6×10^−3^, Figure 2A-D, Movie S3). We obtained similar results when we acutely disrupted integrins by injecting embryos 10 minutes before wounding with 1 mM RGDS (Arg-Gly-Asp-Ser), a peptide that blocks integrin binding to ECM proteins (Naidet et al., 1987; Yasothornsrikul et al., 1997) (Figure S2A-B, Movie S4). RGDS treatment slowed wound closure by 45% (*P* = 9×10^−3^; Figure S2C-D). Our results show that cell-ECM adhesion is necessary for rapid embryonic wound healing.

**Figure 2.**
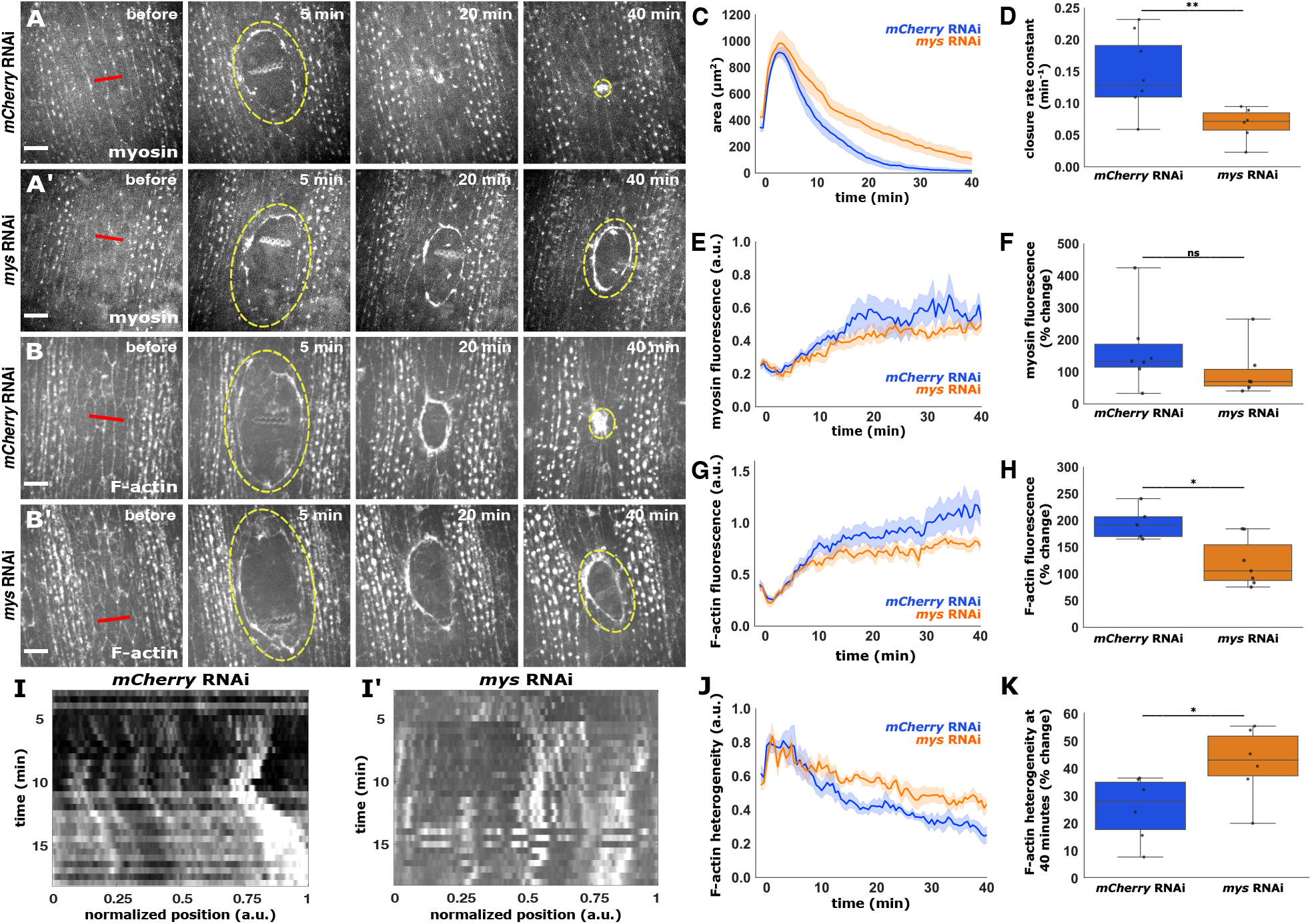
Integrins are necessary for rapid embryonic wound closure and F-actin dynamics at the wound edge. **(A-B**) Wound healing in embryos expressing myosin:GFP (A-A’) or Utrophin:GFP (B-B’) in *mCherry* RNAi (A, B) or *mys* RNAi (A’, B’) embryos. Red lines indicate wound sites. Yellow dashed lines outline the wounds. Anterior left, dorsal up. Bars, 10 μm. **(C-H, J-K)** Wound area over time (C), wound closure rate constant (D), myosin fluorescence at wound edge over time (E), and percent change of myosin fluorescence 40 min after wounding (F) for *mCherry* RNAi (blue, *n* = 8 wounds) and *mys* RNAi embryos (orange, *n* = 6), and F-actin fluorescence at the wound edge over time (G), percent change of F-actin fluorescence 40 min after wounding (H), and F-actin heterogeneity at the wound edge over time (J) and 40 minutes after wounding (K), for *mCherry* RNAi (blue, *n* = 6 wounds) and *mys* RNAi embryos (orange, *n* = 7). **(I**) Kymographs showing F-actin at the wound edge in an *mCherry* RNAi (I) and a *mys* RNAi embryo (I’). (A-B, I) Time after wounding is shown.

### Integrins regulate actin assembly and dynamics at the wound edge

To investigate how integrin disruption affects wound healing, we quantified the formation of the actomyosin cable in embryos expressing myosin II tagged with GFP (Royou et al., 2004) (Figure 2A, Movie S3) or mCherry (Martin et al., 2009) (Figure S2A, Movie S4), or expressing the actin binding domain of Utrophin tagged with GFP as a reporter of filamentous actin (F-actin) (Rauzi et al., 2010) (Figures 2B and S2B, Movies S4 and S5). We did not find a significant difference in myosin accumulation at the wound edge in *mys* RNAi embryos compared to *mCherry* RNAi controls (Figure 2E-F), or in RGDS *vs*. water-treated embryos (Figure S2E-F). In contrast, we measured a significant, 38% decrease in actin levels around the wound 40 minutes after wounding in *mys* RNAi embryos with respect to *mCherry* RNAi controls (*P* = 0.03, Figure 2G-H, Movie S5), and a 40% reduction in RGDS-injected embryos with respect to water-treated controls (*P* = 0.04, Figure S2G-H, Movie S6), indicating that integrins promote actin polarization during embryonic wound closure.

The actin cable around wounds is dynamically remodelled during repair, with the actin distribution starting out heterogeneous and becoming more uniform in response to increasing tension (Kobb et al., 2019; Zulueta-Coarasa and Fernandez-Gonzalez, 2018; Zulueta-Coarasa et al., 2014). To establish if integrins are necessary for the dynamics of actin during wound healing, we measured the heterogeneity of the actin cable, quantified as the ratio between the standard deviation and the mean pixel values. F-actin intensity along the wound edge became more uniform over time in controls, but not in *mys* RNAi embryos: the distribution of actin around the wound 40 minutes after wounding was 91% more heterogeneous in *mys* RNAi embryos compared to controls (*P* = 0.04, Figure 2I-K). Similarly, RGDS-injected embryos displayed an actin distribution at the wound edge 79% more heterogenous than controls (*P =* 0.02, Figure S2I-K). Our results indicate that actin polarization and remodelling at the wound edge requires mechanical engagement of integrins with the ECM. Reduced actin levels at the wound edge could affect the generation of contractile force and delay wound healing (Kobb et al., 2019). Additionally, integrins are necessary for the transmission of tissue-scale forces. In *Drosophila* amnioserosa cells, IBAs act as tethers to the ECM that resist apical deformation by transmitting actin-based forces basally across cells (Goodwin et al., 2016). Disruption of cell-ECM adhesion in the amnioserosa leads to the upregulation of apical force transmission, and changes in actin organization (Goodwin et al., 2017). Our findings suggest that IBAs may be responsible for tension generation, force transmission, or actin organization at the wound edge.

### Integrins are dispensable for force generation at the wound edge

To determine if integrins regulate tension at the wound edge, we used laser ablation to quantify the mechanical properties of the actomyosin cable. We severed the cable when wounds had closed to 50% of their maximum area (Figure 3A-B). The initial recoil velocity after ablation of an actomyosin cable is proportional to the tension sustained (Hutson et al., 2003). We found no difference in the recoil velocity after ablation of the actomyosin cable for control and *mys* RNAi embryos (Figure 3C, Movie S7) or for water and RGDS-treated embryos (Figure S3A-C, Movie S8), suggesting that integrins are dispensable for contractility at the wound edge. To determine if integrins control the viscoelastic properties of the actomyosin cable, we fitted the laser ablation data using a Kelvin-Voigt model to estimate a relaxation time that represents the viscosity-to-elasticity ratio (Kumar et al., 2006; Zulueta-Coarasa and Fernandez-Gonzalez, 2015). The relaxation time was similar for control and *mys* RNAi embryos (Figure 3D), and for water and RGDS-injected embryos (Figure S3D), suggesting that integrins do not control the viscoelasticity of the wound edge.

**Figure 3.**
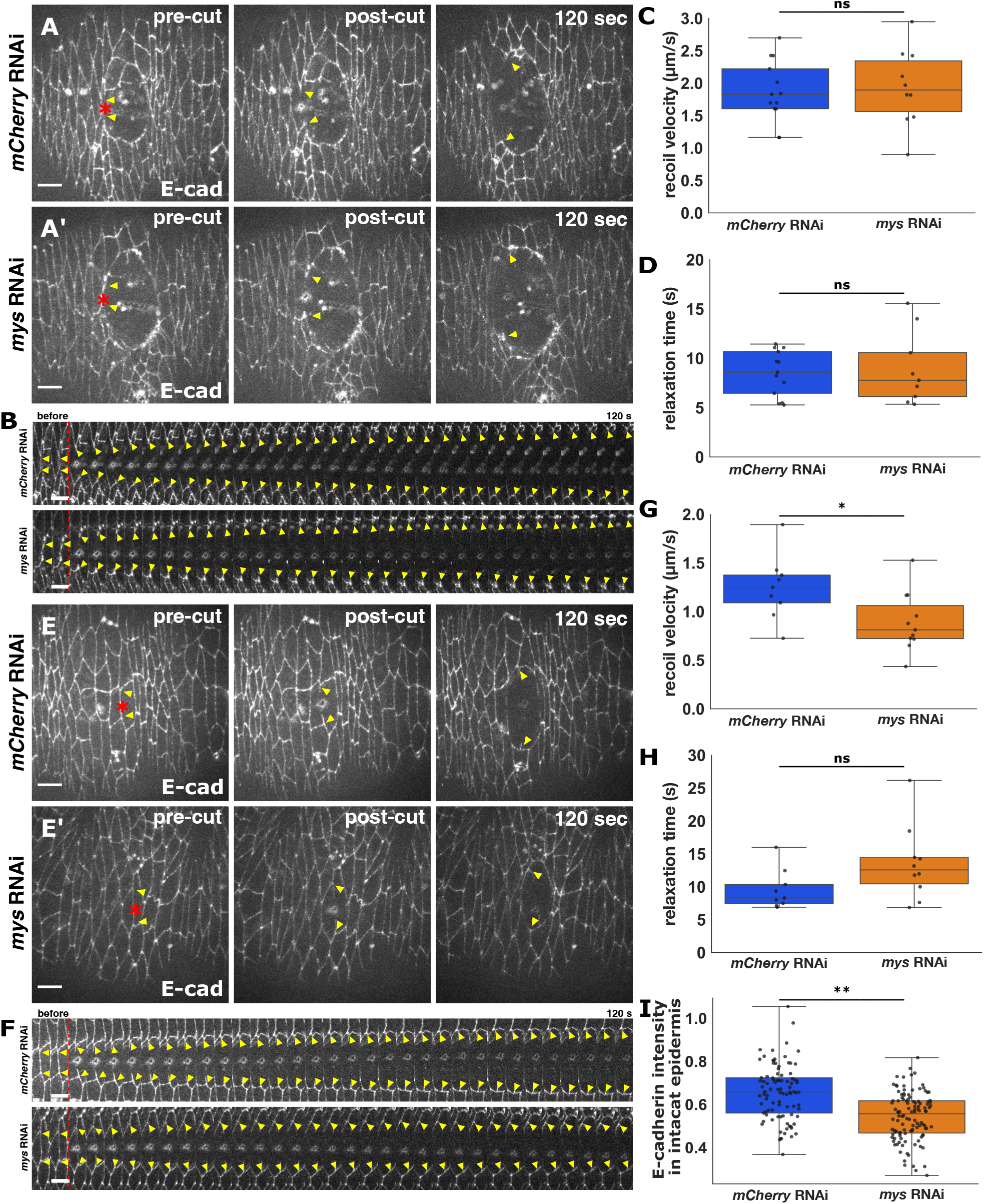
Integrin knock-down does not affect cable tension during embryonic wound repair but reduces baseline epidermal tension. **(A, E)** Laser ablation of a segment of the wound edge (A) or of a cell-cell contact in the intact epidermis (E) in *mCherry* RNAi (A, E) or *mys* RNAi (A’, E’) embryos expressing E-cadherin:tdTomato. Time after wounding is shown. Red asterisks indicate the laser targets. Yellow arrowheads indicate the ends of the severed structure. Anterior left, dorsal up. Bars, 10 μm. **(B, F)** Kymographs showing the structures highlighted by the yellow arrowheads in (A, E), respectively. Bars, 4 s. Red dashed lines indicate the time of ablation. **(C-D, G-I)** Initial recoil velocity (C, G) and relaxation time (D, H) after ablation of a segment of the wound edge (C, D) or a cell-cell junction (G, H) in *mCherry* RNAi (blue, *n* = 13 segments in C, D and 8 junctions in G, H) and *mys* RNAi embryos (orange, *n* = 10 segments in C, D and 10 junctions in G, H). **(I)** Junctional E-cadherin fluorescence in the intact epidermis in *mCherry* RNAi (blue, *n* = 93 cells in 6 embryos) and *mys* RNAi embryos (orange, *n* = 114 cells in 6 embryos).

Coordinated cell movements in epithelia are affected by the mechanical properties of the tissue (Tetley et al., 2019). To determine if the baseline mechanical properties of the epidermis changed when we disrupted integrins, we used laser ablation to sever individual cell-cell junctions in the unwounded epidermis. RNAi-based integrin knock-down reduced the recoil velocity after ablation by 28% (*P* = 0.03, Figure 3E-G, Movie S9), indicating that the baseline epidermal tension decreased when integrins were disrupted. Relaxation times after ablation were not affected by integrin knock-down (Figure 3H). RGDS treatment did not alter neither recoil velocities nor relaxation times (Figure S3G-H, Movie S10). Notably, *mys* RNAi reduced overall E-cadherin levels in the intact epidermis by 17% (*P* = 2×10^−9^, Figure 3E, I), while RGDS treatment did not affect E-cadherin fluorescence (Figure S3I-J), suggesting that differences in E-cadherin levels could explain the reduction in baseline epidermal tension in *mys* RNAi embryos. The different effects of integrin knock-down *vs*. blocking integrin binding to ECM on epidermal tension and E-cadherin levels suggest that integrin expression but not mechanical engagement is necessary for epidermal homeostasis in the *Drosophila* embryo. Alternatively, the prolonged effect of the integrin knock-down by RNAi could lead to changes in E-cadherin expression that the acute inhibition using RGDS does not have time to induce. In addition, integrins can bind ECM proteins in a non-RGD-dependent manner (Naidet et al., 1987), in a process that would not be affected by RGDS treatment. Finally, in *Drosophila* embryos, integrins can adhere to the vitelline membrane (Münster et al., 2019), a protective sac that encloses the embryo, suggesting that alternative binding partners may contribute to integrin signalling during wound healing.

### Integrins regulate cell-cell adhesion dynamics at the wound edge

To investigate if integrins regulate adherens junction remodelling during wound closure, we quantified the dynamics of E-cadherin at bicellular and tricellular junctions at the wound edge (BCJs and TCJs, respectively, Figure 4A-B, Movie S11). In controls, E-cadherin:GFP decreased by 18±8% at BCJs fifteen minutes after wounding (*P =* 4×10^−3^, Figure 4C), and increased by 14±7% at TCJs (*P* = 2×10^−8^, Figure 4D). Integrin knock-down by RNAi did not affect the depletion of E-cadherin from BCJs (15±9%, *P* = 6×10^−5^, Figure 4C), but prevented the accumulation of E-cadherin at TCJs relative to controls (*P* = 0.01), causing a 12±6% reduction of E-cadherin levels at TCJs fifteen minutes after wounding (*P* = 1×10^−6^, Figure 4D). We obtained similar results when we blocked integrin-ECM binding with RGDS (Figure S4A-D, Movie S12). Together, our results reveal a previously unrecognized interplay between cell-cell and cell-ECM adhesion in the collective cell movements that drive embryonic wound closure.

**Figure 4.**
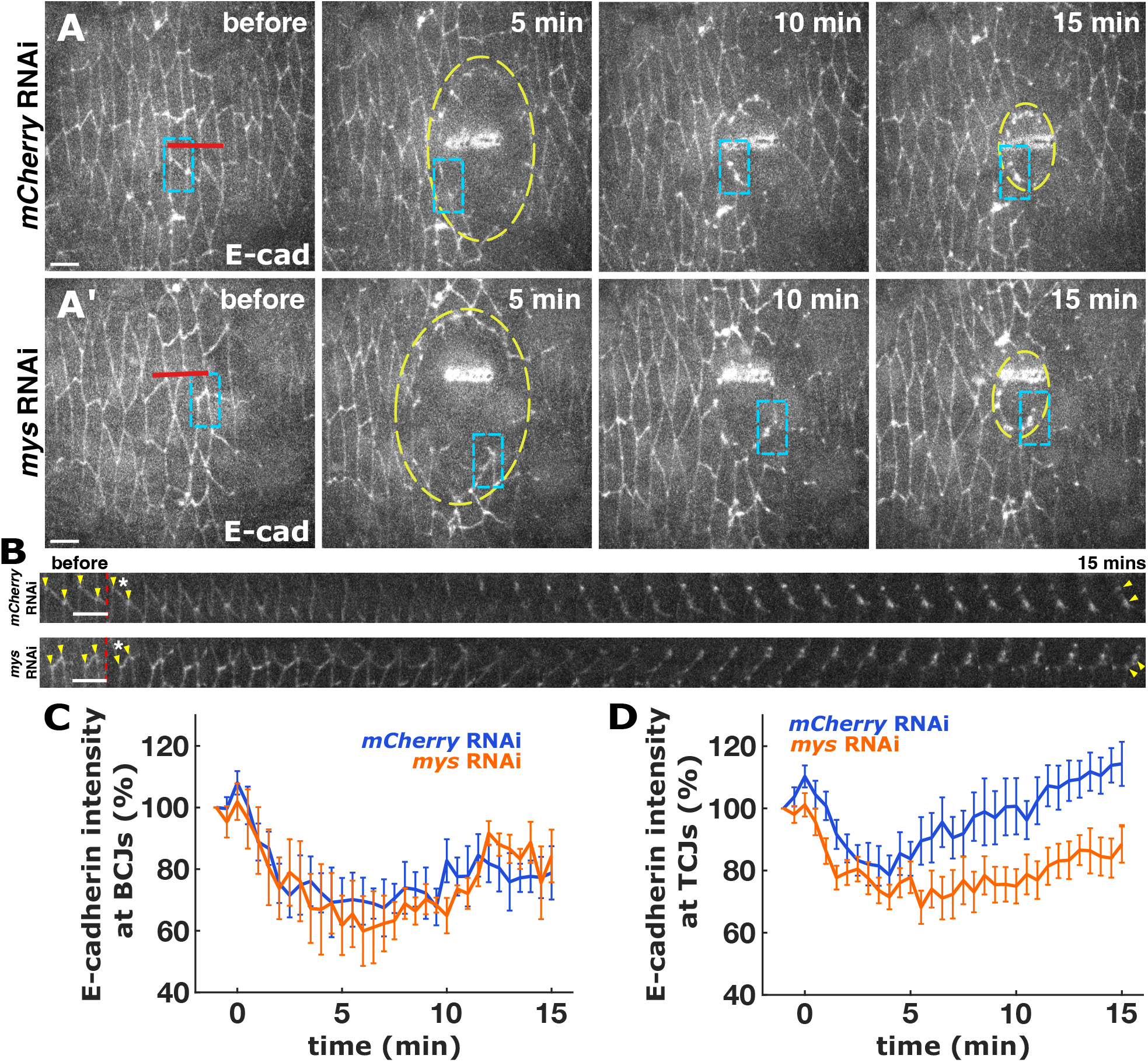
Integrins are necessary for junctional remodelling during embryonic wound repair. **(A)** Wound closure in embryos expressing E-cadherin:GFP and *mCherry* RNAi (A) or *mys* RNAi (A’). Time after wounding is shown. Red lines indicate wound sites. Yellow dashed lines outline the wounds. Cyan indicates the region used for the kymographs in (B). Anterior left, dorsal up. Bars, 10 μm. **(B)** Kymographs showing the junctions highlighted by the cyan boxes in (A). Bars, 30 s. Red dashed lines indicate the time of wounding. Asterisks indicate wound site. Yellow arrowheads indicate TCJs. **(C-D)** Percent change of E-cadherin fluorescence at the wound edge in BCJs (C) and TCJs (D) in embryos expressing *mCherry* RNAi (blue, *n* = 26 BCJs and 41 TCJs in 6 wounds) or *mys* RNAi (orange, n = 21 BCJs and 36 TCJs in 6 wounds).

Crosstalk between cell-cell and cell-ECM adhesions occurs both in tissue culture and *in vivo* and has been proposed to be bidirectional (Bécam et al., 2005; Borghi et al., 2010; Hadjisavva et al., 2022; Jülich et al., 2015). Cadherins and integrins are both transmembrane receptors linked through the actin cytoskeleton, and they share signalling effectors such as non-receptor tyrosine kinases, adaptors, and small GTPases (Weber et al., 2011). Integrins regulate cadherin dependent adhesion during convergent extension movements in *Xenopus* (Marsden and DeSimone, 2003). Similarly, loss of integrin function in *Drosophila* amnioserosa cells leads to defective cadherin localization and reduced stability (Goodwin et al., 2017). Importantly, we find a role for integrins specifically in controlling TCJ remodelling around embryonic wounds. TCJs at the leading edge of collectively migrating cells have been proposed to act as hubs for the nucleation of supracellular actin cables (Kaltschmidt et al., 2002; Matsubayashi et al., 2015). TCJs may also transmit forces across segments of the wound edge that are necessary to stabilize actin around the wound (Kobb et al., 2019). Experiments investigating how integrin disruptions affect E-cadherin trafficking, actin turnover around the wound and the localization and dynamics of different actin regulators will shed light on how integrins contribute to embryonic wound repair.

## MATERIALS AND METHODS

### Fly stocks

Animals were maintained and mated at 18°C or 23°C (room temperature). *Drosophila* strains were kept in plastic vials or bottles on fly food provided by a central kitchen operated by H. Lipshitz. Stage 14-15 embryos (11-12 hours after egg laying) were collected from apple juice-agar plates kept overnight at room temperature (23°C) on collection cages. For immunofluorescence staining, we used *yellow white* or *laminin-A:GFP* flies (Sarov et al., 2016). For live imaging, we used the following markers: *ubi-talin:GFP* (Yuan et al., 2010), *endo-E-cadherin:tdTomato* (Huang et al., 2009), *endo-E-cadherin:GFP (Huang et al*., *2009), sqh-sqh:GFP* (Royou et al., 2004), *sqh-sqh:mCherry* (Martin et al., 2009), and *GFP:UtrophinABD* (Rauzi et al., 2010). We used *UAS-myospheroid* RNAi (Perkins et al., 2015) to knock down integrins, and *UAS-mCherry* RNAi (BL #35785) as a control. UAS constructs were ubiquitously driven with *daughterless*-*Gal4* (Perrin et al., 2003).

### Embryo mounting and drug treatments

*Drosophila* embryos were dechorionated in 50% bleach for 2 minutes and rinsed with water. Embryos were aligned ventral-lateral side up on an apple juice-agar pad and transferred to a coverslip coated with heptane glue. For injections, embryos were dehydrated for 7-10 minutes by placing the coverslip in a plastic container with silica beads (Drierite). After dehydration, embryos were covered with a 1:1 mix of halocarbon oil 27 and 700 (Sigma-Aldrich) (Scepanovic et al., 2021).

Injections were conducted using a Transferman NK2 micromanipulator (Eppendorf) and a PV820 microinjector (World Precision Instruments) coupled to a spinning disk confocal microscope. Embryos were injected with 1 mM RGDS peptide (Tocris Bioscience) in water; or with water as a control. Injections were into the perivitelline space and were followed by an incubation at room temperature for 10 minutes before imaging. Drugs are predicted to be diluted by 50-fold in the perivitelline fluid (Foe and Alberts, 1983).

### Time-lapse imaging

Imaging was performed at room temperature using a Revolution XD spinning disk confocal microscope (Andor Technology) with an iXon Ultra 897 camera (Andor Technology), a 60X oil-immersion lens (NA 1.35; Olympus), and Metamorph software (Molecular Devices). Sixteen-bit Z-stacks were acquired at 0.5 μm steps every 4-30 seconds (11-21 slices per stack). Maximum intensity projections were used for analysis.

### Laser ablations

Wounds on the embryonic epidermis were created using a pulsed Micropoint nitrogen laser (Andor Technology) tuned to 365 nm. Ten laser pulses were delivered in six spots along a 13-μm line to generate a wound. Each embryo was wounded only once. To ensure consistency, we restricted our analyses to wounds larger than 700 μm^2^. For spot laser ablations, ten pulses were delivered at a single point over the course of 670 ms to release tension at the wound margin or in the intact epidermis. Samples were imaged immediately before and 1.72s after spot ablations. After ablation, embryos were imaged every 4 seconds for spot ablations or 30 seconds for wound healing assays.

### Embryo fixation and staining

For immunofluorescent staining, we dechorionated stage 14–15 embryos as above. Embryos were fixed for 20 min in a 1:1 mix of heptane and 37% formaldehyde in phosphate buffer. Embryos were manually devitellinized or popped using 90% ethanol, stained with primary antibodies for 2 hours at room temperature and fluorescently labelled with secondary antibodies incubated for 1 hour at room temperature. Antibodies used were mouse anti-GFP (1:50; DSHB, #DSHB-GFP-12A6), rabbit anti-collagen IV (1:100; Abcam, #ab6586), mouse anti-integrin βPS (0.3 ug/ml; DSHB, #CF.6G11), rat anti-α-catenin (1:20; DSHB, #DCAT-1), rat anti-Discs large (DLG; 1:100; DSHB, #4F3), Alexa 488 goat anti-mouse IgG (1:50; Invitrogen), Alexa 555 goat anti-rat IgG (1:50; Invitrogen), and Alexa 568 goat anti-rabbit IgG (1:500; Invitrogen). Filamentous actin was stained with Alexa 647-conjugated phalloidin (1:500; Invitrogen). Embryos were mounted in ProLong Gold (Molecular Probes) between two coverslips. Collagen and Laminin A stainings were imaged on an Olympus FV3000 laser-scanning confocal microscope with a 60X oil-immersion lens (NA 1.35; Olympus), and FluoView software (Olympus). 16-bit Z-stacks were acquired at 0.5 μm steps (21 slices per stack). Single Z-slices were used for analysis.

### Quantitative image analysis

Image analysis was performed using PyJAMAS (Fernandez-Gonzalez et al., 2022), an image analysis platform developed by our lab, and custom scripts written in Python or in MATLAB (Mathworks). To analyze wound closure phenotypes, we traced wounds using the semi-automated LiveWire method that uses Dijkstra’s optimal path search algorithm to find and trace the brightest path of pixels between two manually selected points (Dijkstra, 1959).

Annotations of TCJs along the wound edge were generated with a semi-automated machine learning method. A support vector machine was trained using PyJAMAS on positive and negative training sets of one hundred 30×30-pixel images each. The positive training set contained images of different TCJs along the wound edge imaged in embryos expressing E-cadherin:tdTomato at various stages of wound closure. The negative training set was created from background regions, bicellular junctions away from the wound, and the auto-fluorescent scars from the wounding laser on the vitelline membrane. The support vector machine was then applied to the images to be annotated. The detected TCJs were filtered by first applying non-maximum suppression to remove low confidence candidates, followed by removing any detections not intersecting the wound edge. TCJs were automatically tracked in time based on the closest Euclidean distance.

We quantified fluorescence at the wound margin as the mean pixel value under a 0.6-μm-wide mask over the wound edge generated by the Livewire algorithm. We used the image mode as the value for background subtraction, and we corrected for photobleaching by dividing by the mean image intensity at each time point. We quantified the fluorescent markers from the pixel values at the wound edge in maximum intensity projections. We measured the percent change of fluorescence as:

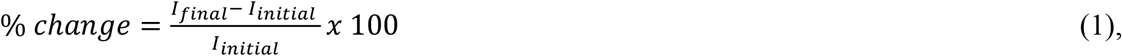

where *I*_*initial*_ is the mean intensity at the wound edge across the first two time points after wounding, when fluorescence at the wound edge reaches its minimum, and *I*_*final*_ is the mean of the last two time points. The basal-to-apical ratio of collagen and Laminin A was measured by tracing 9-16-μm-long segments of the basal and apical surfaces of the epidermis from image cross-sections using the LiveWire algorithm and computing the mean pixel value along each segment. To measure the effect of *mys* RNAi, we quantified integrin βPS fluorescence staining as the mean of a 200 × 200-pixel region on maximum intensity projections of 3-μm stacks of the epidermis. We calculated actin heterogeneity as the ratio between the standard deviation and the mean of the pixel values around the wound.

To create kymographs of the wound edge, annotations from the first 15 minutes after wounding were circularly shifted until the start points were aligned. The intensity signal along the wound edge was interpolated to obtain 1500 equally spaced points. The intensity signal was corrected for background intensity and photobleaching before being smoothed with a Gaussian kernel with a size of 5 pixels (1.33 μm). Each row of the kymograph displays the intensity of the wound edge at a single time point. Three-dimensional reconstructions of E-cadherin and Talin were created using the ClearVolume plugin (Royer et al., 2015) for Fiji (Schindelin et al., 2012).

To measure retraction velocity after laser ablation, the positions of the two tricellular vertices connected by the ablated structure (wound edge segment or cell-cell junction) were manually tracked in PyJAMAS. We quantified the retraction velocity of TCJs after ablation as a proxy for mechanical tension (Hutson et al., 2003; Zulueta-Coarasa and Fernandez-Gonzalez, 2015). We used a Kelvin-Voigt mechanical equivalent circuit, which models junctions as a combination of a spring and a dashpot configured in parallel, to estimate the viscosity-to-elasticity ratio (Zulueta-Coarasa and Fernandez-Gonzalez, 2015).

### Statistical analysis

To measure the significance of changes across two unpaired samples, we used a non-parametric Mann-Whitney test. To measure the significance of change of a single sample relative to zero, we used a one-sample Wilcoxon signed-rank test. In box plots, boxes show the quartiles, error bars indicate the range, and the grey line is the median. In time-course plots, error bars represent the standard error of the mean. ns – not significant, * *P* < 0.05, ** *P* < 0.01.

## ACKNOWLEDGEMENTS

We thank Guy Tanentzapf for reagents. Flybase provided important information for this study. We are grateful to Ana Maria do Carmo and Gordana Scepanovic for comments on the manuscript. KER was supported by postdoctoral fellowships from the Canadian Institutes of Health Research and the Ted Rogers Centre for Heart Research. This work was funded by grants to RFG from the Canadian Institutes of Health Research (156279), the Natural Sciences and Engineering Research Council of Canada (418438-13), and the University of Toronto Translational Biology and Engineering and XSeed Programs. RFG is the Canada Research Chair in Quantitative Cell Biology and Morphogenesis.

