## Supplemental Figures and Movie Legends for "Integrin-based adhesions promote cell-cell junction remodelling and cytoskeletal rearrangements to drive embryonic wound healing"

### SUPPLEMENTARY FIGURES

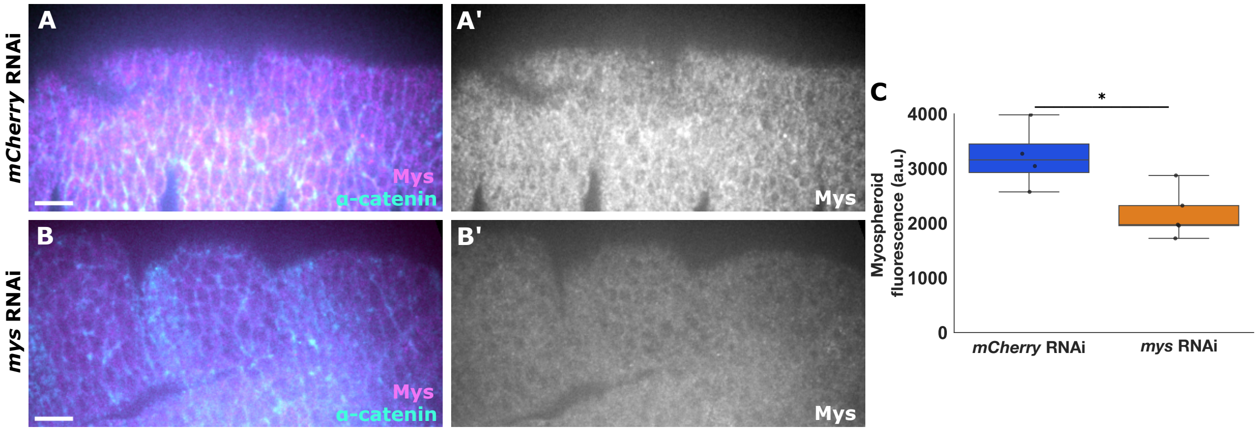


**Figure S1. *mys* RNAi reduces Mys levels in the embryonic epidermis. (A-B)** Maximum intensity projection of 3 μm stacks of the epidermis of *Drosophila* embryos expressing *mCherry* RNAi (A) or *mys* RNAi (B) and stained with antibodies against *mys* (magenta, grayscale in A'-B') and α-catenin (cyan). Anterior left, dorsal up. Bars, 10 μm. **(C)** Myospheroid fluorescence in the epidermis of embryos expressing *mCherry* RNAi ($n$ = 4) or *mys* RNAi ($n$ = 5).

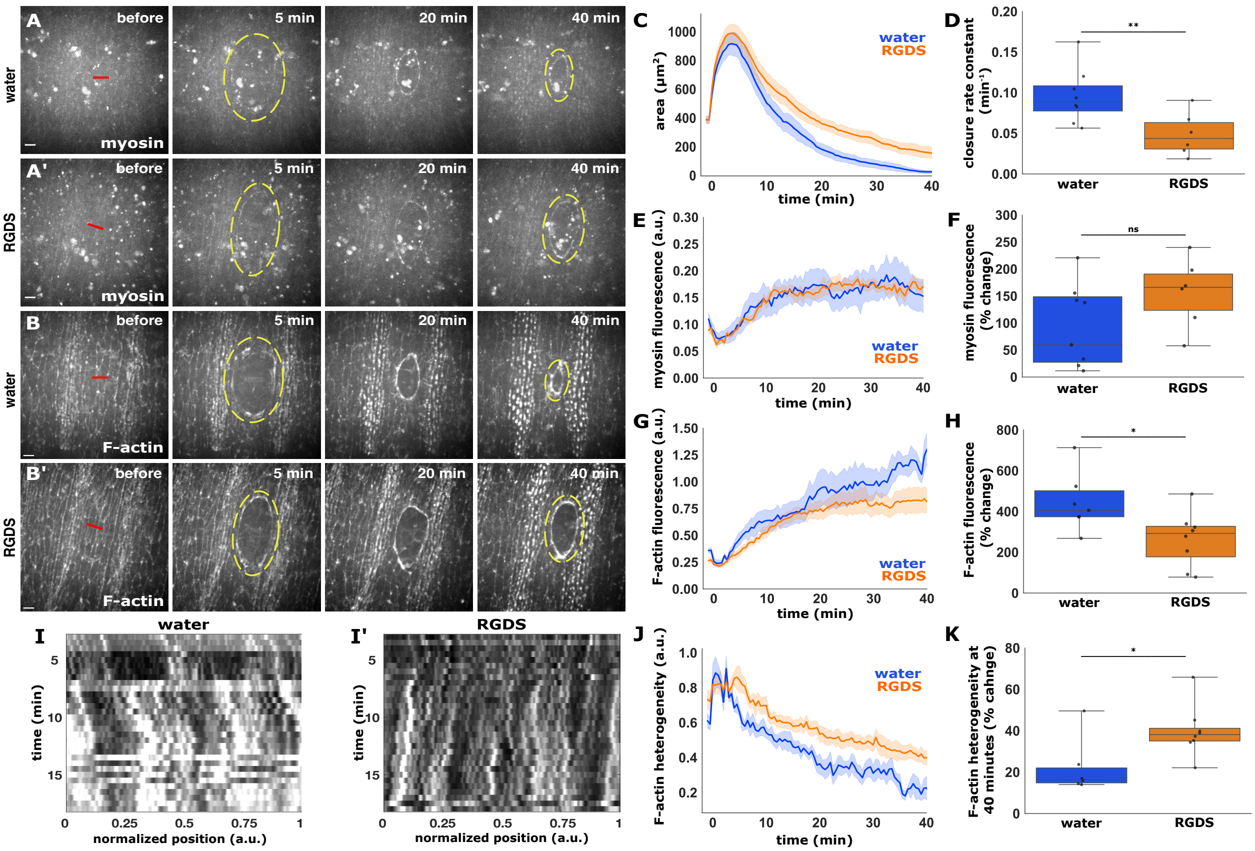


**Figure S2. Blocking integrin binding disrupts wound closure and F-actin dynamics. (A-B)** Wound closure in embryos expressing myosin:mCherry (A-A') or Utrophin:GFP (B-B') and injected with water (A, B) or RGDS (A', B'). Anterior left, dorsal up. Red lines indicate wound sites. Yellow dashed lines outline the wounds. Bars, 10 µm. **(C-H)** Wound area over time (C), wound closure rate constant (D), myosin fluorescence at wound edge over time (E), percent change of myosin fluorescence 40 min after wounding (F), F-actin fluorescence at the wound edge over time (G), and percent change of F-actin fluorescence 40 min after wounding (H) for water-injected (blue, *n* = 8 wounds in C-F and 6 wounds in G-H) and RGDS-injected embryos (orange, *n* = 6 wounds in C-F and 8 wounds in G-H). **(I)** Kymographs showing F-actin at the wound edge in a water-injected (I) and a RGDS-injected embryo (I'). **(J-K)** F-actin heterogeneity at the wound edge over time (J) and 40 minutes after wounding (K), for water-injected (blue, *n* = 6 wounds) and RGDS-injected (orange, *n* = 8) embryos. (A-B, I) Time after wounding is shown.

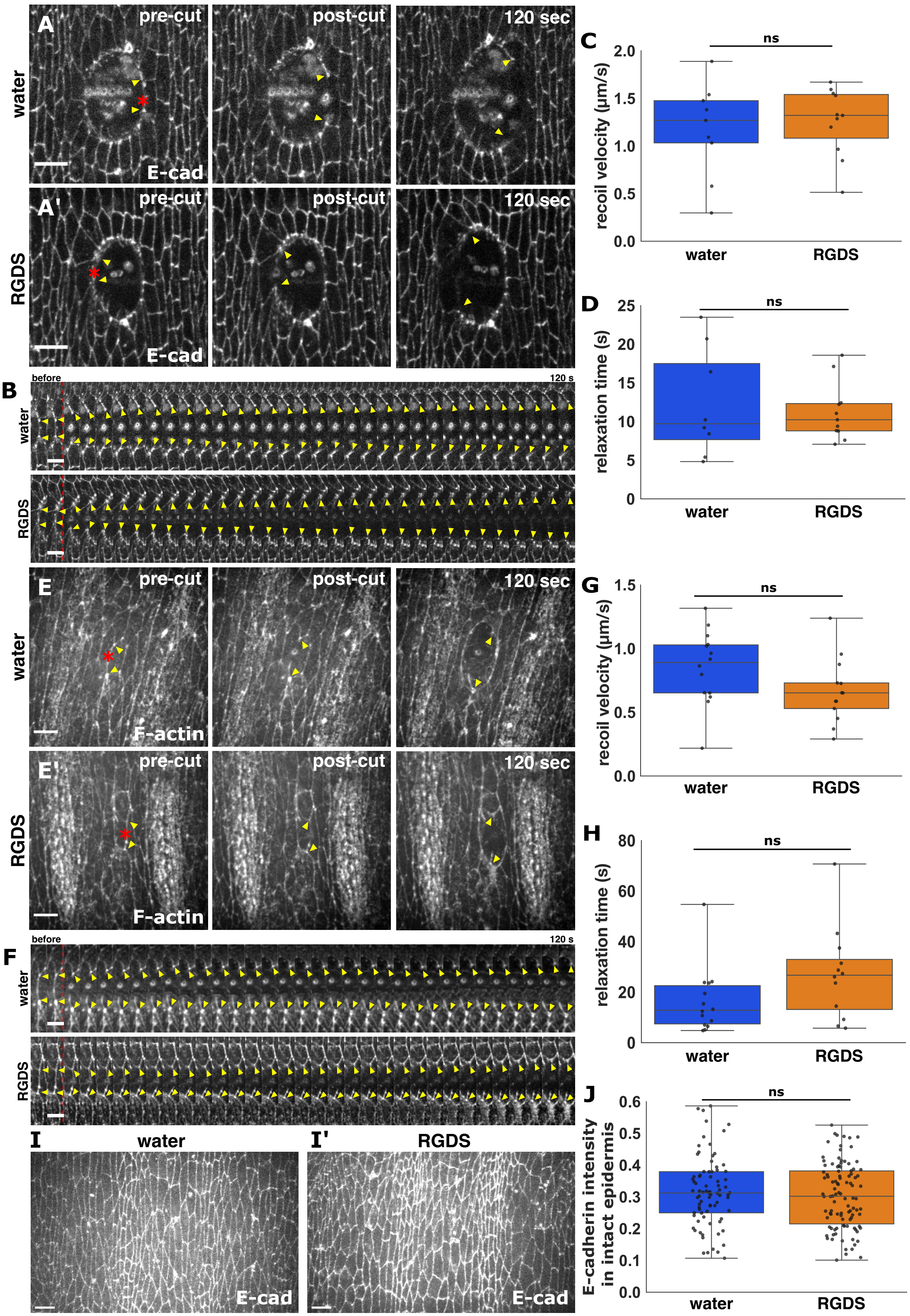

A
pre-cut
post-cut
120 sec
water
E-cad
A'
RGDS
B
before
120 s
C
ns
recoil velocity (μm/s)
2.0
1.5
1.0
0.5
0.0
water
RGDS
D
relaxation time (s)
25
20
15
10
5
0
E
F-actin
E'
F
G
1.5
H
80
60
40
20
I
I'
J
E-cadherin intensity in intact epidermis
0.6
0.5
0.4
0.3
0.2
0.1

**Figure S3. Blocking integrins does not affect cell mechanics in the *Drosophila* embryonic epidermis. (A, E)** Laser ablation of a segment of the wound edge (A) or a cell-cell contact in the intact epidermis (E) in embryos treated with water (A, E) or RGDS (A', E') and expressing E-cadherin:tdTomato. Time after wounding is shown. Red asterisks indicate the laser targets. Yellow arrowheads indicate the ends of the severed structure. Anterior left, dorsal up. Bars, 10 μm. **(B, F)** Kymographs showing the structures highlighted by the yellow arrowheads in (A, E), respectively. Bars, 4 s. Red dashed lines indicate the time of ablation. **(C-D, G-H)** Initial recoil velocity (C, G) and relaxation time (D, H) after ablation of a segment of the wound edge (C, D) or a cell-cell junction (G, H) in embryos injected with water (blue, *n* = 9 segments in C, D and 7 junctions in G, H) or RGDS (orange, *n* = 11 segments in C, D and 8 junctions in G, H). **(I)** Epidermal cells in embryos expressing E-cadherin:tdTomato and injected with water (I) or RGDS (I'). (**J**) Junctional E-cadherin fluorescence in water (blue, *n* = 80 cells in 7 embryos) or RGDS-injected embryos (orange, *n* = 102 cells in 10 embryos).

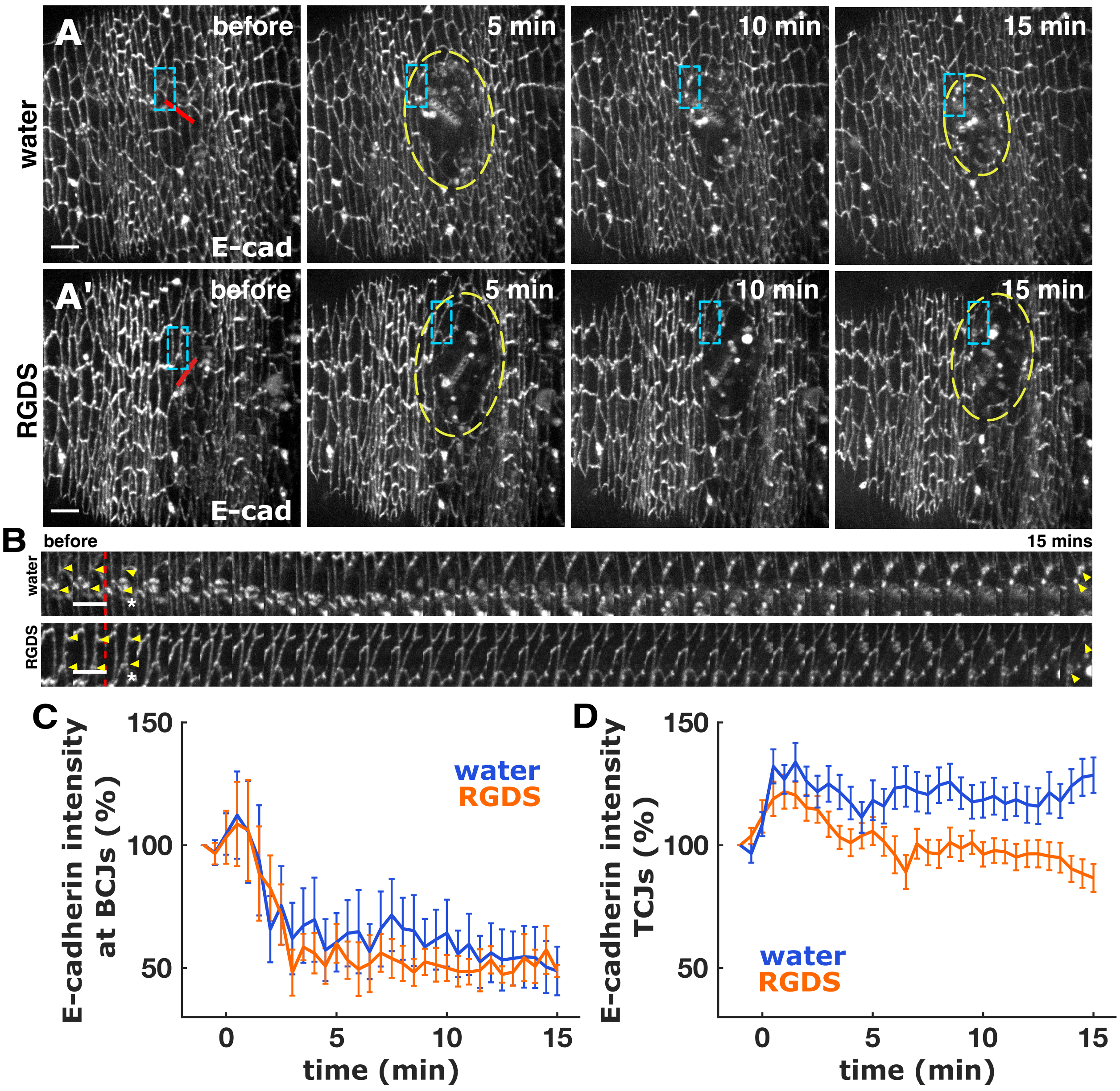


**Figure S4. Integrin-based adhesion is necessary for junctional remodelling during embryonic wound repair. (A)** Wound closure in embryos expressing E-cadherin:GFP and treated with water (A) or RGDS (A'). Time after wounding is shown. Red lines indicate the wound sites. Yellow dashed lines outline the wounds. Cyan indicates the region used for the kymographs in (B). Anterior left, dorsal up. Bars, 10 µm. **(B)** Kymographs showing the junctions highlighted by the cyan boxes in (A). Bars, 30 s. Red dashed lines indicate the time of wounding. Asterisks indicate wound sites. Yellow arrowheads indicate TCJs. **(C-D)** Percent change of E-cadherin fluorescence at the wound edge in BCJs (C) and TCJs (D) in embryos injected with water (blue, *n* = 25 BCJs and 86 TCJs in 7 wounds) or RGDS (orange, *n* = 27 BCJs and 93 TCJs in 8 wounds).

### SUPPLEMENTARY MOVIE LEGENDS

**Movie S1. Talin accumulates at the wound edge.** Wound healing in an embryo expressing Talin:GFP. Images were acquired every 30 s. Time after wounding is shown. Anterior left, dorsal up. Bar, 10 μm.

**Movie S2. Talin colocalizes with E-cadherin at the wound edge.** Wound edge 15 minutes after wounding in an embryo expressing Talin:GFP (green) and E-cadherin:tdTomato (magenta). Anterior left, dorsal up.

**Movie S3. Integrins are necessary for rapid wound closure but dispensable for myosin polarization to the wound edge.** Wound healing in embryos expressing myosin:GFP and *mCherry* RNAi (left) or *mys* RNAi (right). Images were acquired every 30 s. Time after wounding is shown. Anterior left, dorsal up. Bar, 10 μm.

**Movie S4. Blocking integrin signalling disrupts wound repair but does not affect myosin polarization to the wound edge.** Wound healing in embryos expressing myosin:mCherry and injected with water (left) or RGDS (right). Images were acquired every 30 s. Time after wounding is shown. Anterior left, dorsal up. Bar, 10 μm.

**Movie S5. Integrins are necessary for actin accumulation at the wound edge**. Wound healing in embryos expressing GFP:UtrophinABD and *mCherry* RNAi (left) or *mys* RNAi (right). Images were acquired every 30 s. Time after wounding is shown. Anterior left, dorsal up. Bar, 10 μm.

**Movie S6. Blocking integrins disrupts actin accumulation at the wound edge.** Wound healing in embryos expressing GFP:UtrophinABD and injected with water (left) or RGDS (right). Images were acquired every 30 s. Time after wounding is shown. Anterior left, dorsal up. Bar, 10 μm.

**Movie S7. Integrins are not necessary to generate tension at the wound edge.** Laser ablation of the wound edge in embryos expressing E-cadherin:GFP and *mCherry* RNAi (left) or *mys* RNAi (right). Images were acquired every 4 s. Time after wounding is shown. Anterior left, dorsal up. Bar, 10 μm.

**Movie S8. Blocking integrins does not affect tension around the wound.** Laser ablation of the wound edge in embryos expressing E-cadherin:GFP and injected with water (left) or RGDS (right). Images were acquired every 4 s. Time after wounding is shown. Anterior left, dorsal up. Bar, 10 μm.

**Movie S9. Integrins are necessary to maintain baseline epidermal tension.** Laser ablation of a cell junction in the intact epidermis of embryos expressing E-cadherin:GFP, and *mCherry* RNAi (left) or *mys* RNAi (right). Images were acquired every 4 s. Time after wounding is shown. Anterior left, dorsal up. Bar, 10 μm.

**Movie S10. Blocking integrins does not affect epidermal tension.** Laser ablation on a cell junction in the intact epidermis of embryos expressing GFP:UtrophinABD and injected with water (left) or RGDS (right). Images were acquired every 4 s. Time after wounding is shown. Anterior left, dorsal up. Bar, 10 μm.

**Movie S11. Integrins are necessary for E-cadherin rearrangement during wound closure.** Wound healing in embryos expressing endo-Ecadherin:GFP and *mCherry* RNAi (left) or *mys* RNAi (right). Images were acquired every 30 s. Time after wounding is shown. Anterior left, dorsal up. Bar, 10 μm.

**Movie S12. Blocking integrins disrupts E-cadherin remodelling during embryonic wound repair.** Wound healing in embryos expressing endo-Ecadherin:GFP and injected with water (left) or RGDS (right). Images were acquired every 30 s. Time after wounding is shown. Anterior left, dorsal up. Bar, 10 μm.
